# Transcriptome profiles in brains of mice heterozygous for a DYT1 dystonia-associated mutation in the endogenous *Tor1a* gene

**DOI:** 10.1101/825505

**Authors:** Sara B. Mitchell, Michael S. Chimenti, Hiroyuki Kawano, Tsun Ming Tom Yuen, Ashley E. Sjurson, Sadahiro Iwabuchi, Kevin L Knudtson, Thomas B Bair, Diana Kolbe, N. Charles Harata

## Abstract

In patients with the brain disorder dystonia, body movement is severely affected – with involuntary muscle contractions and abnormal postures, causing extensive deterioration of the patient’s quality of life. The most common inherited form of this disorder is DYT1 dystonia, which is caused by a mutation in *TOR1A* gene and autosomal dominant. The molecular mechanisms that underlie the effects of the *TOR1A* mutation on brain function remain unclear. To understand these, we examined the gene expression profiles (transcriptome) in four brain regions (cerebral cortex, hippocampus, striatum and cerebellum) in a mouse model, the heterozygous ΔE-torsinA knock-in mice which genetically reproduce the mutation in DYT1 dystonia. The samples were obtained at 2 to 3 weeks of age, a period during which synaptic abnormalities have been reported. Pairwise comparisons of brain regions revealed differential gene expression irrespective of genotype. A comparison of heterozygous to wild-type mice failed to reveal genotype-dependent differences in gene expression in any of the four brain regions when examined individually. However, genotype-dependent differences became apparent when the information for all brain regions was combined. These results suggest that any changes in the transcriptome within a brain region were subtle at this developmental stage, but that statistically significant changes occur across all brain regions. Such changes in the transcriptome, although subtle in degree, could underlie the processes that give rise to DYT1 dystonia.

## 1. Introduction

DYT1 dystonia is defined as “early-onset generalized isolated dystonia” (Albanese et al., 2013). It is characterized by involuntary muscle contractions and abnormal postures, and caused by a mutation in the *TOR1A* gene (c.904_906delGAG/c.907_909delGAG; p.Glu302del/p.Glu303del) that results in deletion of a glutamic acid residue from the protein product, torsinA (ΔE-torsinA) (Bragg et al., 2011; Breakefield et al., 2008; Geyer and Bressman, 2006; Ozelius et al., 1997; Tanabe et al., 2009). Patients with this autosomal dominant disorder are heterozygous for the mutation (*TOR1A*^+/ΔE^). TorsinA belongs to the <u>A</u>TPases <u>a</u>ssociated with various cellular <u>a</u>ctivities (AAA^+^) protein superfamily (Hanson and Whiteheart, 2005). Its ATPase activity is regulated by the interacting partners of torsinA (Brown et al., 2014; Sosa et al., 2014; Zhao et al., 2013), and is thought to contribute to diverse cellular processes including the unfolding of proteins during their degradation, the disassembly of protein complexes and aggregates, and the regulation of cellular responses to stress (Beauvais et al., 2016; Beauvais et al., 2018; Gonzalez-Alegre, 2019; Hanson and Whiteheart, 2005; Nery et al., 2011; Rittiner et al., 2016). Although some specific biochemical properties of the torsinA protein have been identified, the affected pathways may be diverse, and they could be cell- and region-dependent. Thus the molecular mechanisms whereby the *TOR1A* mutation contributes to neuronal dysfunction in the brain (Kakazu et al., 2012a; Kakazu et al., 2012b; Martella et al., 2014; Song et al., 2012) remain elusive.

Transcriptome analysis can be an effective way to obtain insight into the molecular pathophysiology of diseases. This approach reveals changes in the expression levels of a large number of mRNAs, and it does so in an unbiased manner. Thus, it is not limited to the analysis of a few pre-selected mRNA targets, and it offers the advantage of integrating primary and secondary consequences of the initial cause, e.g. gene mutation, and thus characterizing the overall pathological condition. Any changes that are detected represent a readout of a given condition, and if changes are large, the analysis would make it possible to semi-quantitatively assess which brain regions and gene networks are most affected.

A number of previous studies of DYT1 dystonia included the analysis of transcriptome profiles (**Table 1**). These reports suggested either that there was no change or that the changes were minimal or modest in terms of the numbers of differentially expressed genes. However, the experimental systems assessed in these studies were diverse. Samples ranged from cultured human embryonic kidney (HEK) cells (Baptista et al., 2003), cultured neuron-like pheochromocytoma (PC) cells of rat (Martin et al., 2009), freshly-drawn peripheral blood cells of humans (Walter et al., 2010) to mouse brain (Beauvais et al., 2018; Grundmann et al., 2008). The nature of introducing gene mutation differed across studies, with some using overexpression of mutant, human *TOR1A* gene (Baptista et al., 2003; Grundmann et al., 2008; Martin et al., 2009) and others using mutated allele of the endogenous gene as in the patients (Beauvais et al., 2018; Walter et al., 2010). The only study to evaluate the transcriptome in human patients used blood cells (Walter et al., 2010), and it is likely that the transcription profiles of these cells differ markedly from those for their brain counterparts. In summary, it remains unclear how the transcriptome profiles in brain cells, in particular, how the profiles in the brain regions most related to dystonia pathophysiology, are affected in the context of DYT1 dystonia.

**Table 1.** Transcriptome profiles in the mammalian systems associated with DYT1 dystonia.

| **Mutations or nature of gene mutations** | **Host species (age)** | **Preparation** | **Method (number of samples per comparison)** | **Findings** | **Reference** |
| --- | --- | --- | --- | --- | --- |
| *TOR1A*<sup>ΔE</sup> or *TOR1A*<sup>+</sup> overexpression | human | cultured HEK cells | microarray (n = 2, with 4 cell lines pooled into 2 groups) | #DEx = 39 (1.5 ≤ FC ≤ 2.5), up- or down-regulated in the same direction in *TOR1A*<sup>ΔE</sup> or *TOR1A*<sup>+</sup> vs. vector-only control | (Baptista et al., 2003) |
| *TOR1A*<sup>ΔE</sup> or *TOR1A*<sup>+</sup> overexpression | mouse (3 months) | brain (striatum) | microarray (n = 3 mice) | #DEx = 34 (2.0 ≤ FC ≤ 3.5) in *TOR1A*<sup>ΔE</sup> vs. *TOR1A*<sup>+</sup>, but 502 (2.0 ≤ FC ≤ 17.1) in overexpression of *TOR1A*<sup>ΔE</sup> or 338 (2.0 ≤ FC ≤ 20.6) in overexpression of *TOR1A*<sup>+</sup> alone vs. non-transgenic control mice | (Grundmann et al., 2008) |
| *TOR1A*<sup>ΔE</sup> or *TOR1A*<sup>+</sup> overexpression | rat | cultured PC6-3 cells | microarray (n = 3) | no changes (FC < 1.5) in *TOR1A*<sup>ΔE</sup> or *TOR1A*<sup>+</sup> vs. their non-overexpressing controls (i.e. no treatment with doxycycline) | (Martin et al., 2009) |
| *TOR1A*<sup>+ / ΔE</sup> | human patients & non-manifesting carriers (46.2 & 55.4 years, mean values) | peripheral blood cells | microarray (n = 15 patients, 15 non-manifesting, 9 control subjects) | #DEx = 256 (2.0 ≤ FC ≤ 3.9) in patients vs. non-manifesting, 1509 (2.1 ≤ FC ≤ 15.0) in patients vs. controls, 823 (2.1 ≤ FC ≤ 5.0) in non-manifesting vs. control subjects | (Walter et al., 2010) |
| *Tor1a*<sup>+ / ΔE</sup>, *Tor1a*<sup>ΔE / ΔE</sup> knock-in | mouse (E18) | brain (striatum) | RNA-seq (n = 4) | #DEx = 824 *Tor1a*<sup>+ / ΔE</sup>, 1599 *Tor1a*<sup>ΔE / ΔE</sup>, 98 overlap (intersection) vs. wild-type controls | (Beauvais et al., 2018) |
| *Tor1a*<sup>+ / ΔE</sup> knock-in | mouse (2-3 weeks) | brain (striatum, cerebellum, cerebral cortex, hippocampus) | microarray (n = 9 mice / brain region / genotype) (combined n = 36 samples / genotype) | no changes in individual brain regions except for a single gene (FC = 1.22), but #DEx = 301 (mostly 0.81 < FC < 1.32) in *Tor1a*<sup>+ / ΔE</sup> vs. wild-type controls in the combined samples | current work |

#DEx, number of differentially expressed genes. FC, fold change in a linear scale (i.e. not a logarithmic scale). FC values represent the results of analyses, not the threshold for detecting differentially expressed genes.

In developing our strategy to evaluate transcriptome changes in animal models of DYT1 dystonia, we considered three key issues: genotype, brain regions and developmental stage. The first among these was selection of the most appropriate *genotype* for the analysis. DYT1 dystonia patients are heterozygous for the ΔE-torsinA mutation (*TOR1A*^+/ΔE^). In this sense, the best animal model to date appears to be the heterozygous ΔE-torsinA knock-in mouse model (*Tor1a*^+/ΔE^ or HET) (Beauvais et al., 2018; Dang et al., 2005; Goodchild et al., 2005). It accurately replicates the human condition in that the mice are heterozygous for the mutant allele of endogenous *Tor1a* gene, rather than overexpressing a mutant form of heterologous transgenes or lacking the endogenous wild-type (WT) gene. This is particularly important, given that overexpression of either a mutant form or the WT human *TOR1A* gene has been reported to change the transcriptome in the WT host environment (Baptista et al., 2003; Grundmann et al., 2008), rendering interpretation of overexpression studies difficult (**Table 1**).

Furthermore, both the overexpression and knock-out of a WT gene leads to the formation of morphologically abnormal vesicles between the inner and outer nuclear membranes (“blebbing”), a phenomenon that is not observed in cells of *Tor1a*^+/ΔE^ mice (Kim et al., 2010). Importantly, torsinA proteins are neither overexpressed nor eliminated in human patients; in fact, they are expressed at levels similar to (Goodchild and Dauer, 2004) or slightly lower than (Cao et al., 2010; Goodchild et al., 2005; Yokoi et al., 2012) those in control subjects.

The second point we considered was selection of the most appropriate *brain regions* to analyze. The striatum, which is relevant to motor control, has long been thought to be the major brain region responsible for dystonia pathophysiology (Neychev et al., 2011; Pisani et al., 2007; Tanabe et al., 2009). Conditional knock-out of the *Tor1a* gene in the mouse striatum leads to motor deficits (Yokoi et al., 2011). Also, in *Tor1a*^+/ΔE^ mice, both the morphology of striatal neurons (Song et al., 2013) and synaptic transmission (Martella et al., 2014; Rittiner et al., 2016; Sciamanna et al., 2014) are abnormal. Similar changes in synaptic transmission (Martella et al., 2009) and biochemical signaling (D’Angelo et al., 2017; Sciamanna et al., 2014) were observed in the striatum of transgenic mice that overexpress ΔE-torsinA. Because of the recognition of this important role of the striatum, transcriptome studies of brains have focused solely on this structure (Beauvais et al., 2018; Grundmann et al., 2008) (**Table 1**). However, DYT1 dystonia is increasingly becoming recognized as a disorder of neural networks (DeSimone et al., 2016; Neychev et al., 2011; Prudente et al., 2014; Tanabe et al., 2009; Vo et al., 2015). It suggests both that multiple brains regions are involved in the pathogenesis, and that distinct brain regions might contribute to different aspects of the dystonia pathophysiology. From the viewpoint of transcriptome, it is possible that the potential transcriptome changes are unique in distinct brain regions. Conversely, it is possible that transcriptome changes are similar throughout the brain, because *Tor1a* gene is expressed universally, though at different levels (Allen_Mouse_Brain_Atlas, 2009; Augood et al., 1999; Augood et al., 1998). These alternative possibilities have not been addressed. In this work, we will address them by evaluating multiple brain regions. As described below, other brain regions that might be of particular relevance to DYT1 dystonia include the cerebral cortex, the cerebellum and the hippocampus.

Evidence of a role for the cerebral cortex includes the fact that cortical excitability is enhanced in patients with DYT1 dystonia (Carbon et al., 2004; Edwards et al., 2006; Edwards et al., 2003). In the mouse, conditional knock-out of *Tor1a* in the cerebral cortex leads to motor deficits (Yokoi et al., 2008). Moreover, cerebral cortical pyramidal cells serve as presynaptic neurons for cortico-striatal glutamatergic synaptic transmission, and the efficiency of transmission at this synapse is enhanced in *Tor1a*^+/ΔE^ mice (Martella et al., 2014; Rittiner et al., 2016), as well as in transgenic animals overexpressing ΔE-torsinA pan-neuronally (Grundmann et al., 2012; Martella et al., 2009; Sciamanna et al., 2012; Sciamanna et al., 2011).

The cerebellum has likewise been implicated in dystonia based on multiple observations (Shakkottai et al., 2017; Tewari et al., 2017). Firstly, both the *Tor1a* mRNA (Allen_Mouse_Brain_Atlas, 2009; Augood et al., 1999; Augood et al., 1998) and the torsinA protein (Shashidharan et al., 2000; Walker et al., 2001) are expressed abundantly in the cerebellum. Secondly, cerebellar abnormalities at the network level are observed in mouse models with *Tor1a* gene mutations. For instance, the cerebello-thalamo-cortical pathway is thinned in *Tor1a*^+/ΔE^ mice and the metabolic activity in the sensorimotor cerebral cortex is reduced correspondingly (Ulug et al., 2011). In transgenic mice overexpressing a human ΔE-torsinA, the metabolic activity was increased in both the cerebellar cortex and the inferior olive that sends climbing-fiber input to the cerebellar Purkinje neurons (Zhao et al., 2011). These results imply that cerebellar changes in neuronal connection and metabolism might be involved in the pathophysiology of DYT1 dystonia. Thirdly, cerebellar abnormalities at the cell level are induced by *Tor1a* gene mutations. Cerebellar Purkinje neurons showed few dendritic spines (Song et al., 2014; Zhang et al., 2011), and shortened (Zhang et al., 2011) or thinned dendrites (Song et al., 2014) in *Tor1a*^+/ΔE^ mice and mice with conditional knock-out of *Tor1a* gene. Cerebellar synaptogenesis during early postnatal period was also compromised in mice with *Tor1a* gene mutations (Vanni et al., 2015). Fourthly, acute *Tor1a* knock-down in the cerebellum induced dystonic movements in mice (Fremont et al., 2017).

Evidence for involvement of the hippocampus in dystonia includes the fact that it is one of the regions in which *Tor1a* mRNA (Allen_Mouse_Brain_Atlas, 2009; Augood et al., 1999; Augood et al., 1998) and torsinA protein (Shashidharan et al., 2000; Walker et al., 2001) are especially abundant. In addition, torsinA expression in hippocampal neurons is upregulated as a consequence of ischemia, suggesting that torsinA plays a role in neuronal stress within this region (Zhao et al., 2008). Hippocampal cells also express proteins that are associated with other forms of dystonia, for example, ε-sarcoglycan (Ritz et al., 2011). Furthermore, the hippocampus is one of the regions that are most highly activated in rodents after dystonia-associated behaviors are triggered by the administration of L-type Ca^2+^ channel activators (Jinnah et al., 2003). Although the hippocampus is classically considered as unrelated to movement control, it can play a role in non-motor signs of the mutation carriers.

The third point considered in our experimental design was the most relevant *developmental stage* to test. One key aspect taken into account was the fact that the expression of torsinA in mouse and human brains increases during embryonic period and reaches a peak neonatally (Shashidharan et al., 2000; Siegert et al., 2005; Vasudevan et al., 2006). The embryonic transcriptome was evaluated in the striatum of *Tor1a*^+/ΔE^ mice (Beauvais et al., 2018). After this phase, the torsinA expression decreases rapidly during a postnatal 2-to-3-week period (Puglisi et al., 2013; Tanabe et al., 2016). This early postnatal time is the critical period during which *Tor1a* mutations lead to abnormalities in: synaptogenesis in the cerebellum (Vanni et al., 2015); the plasticity of synaptic transmission at the cortico-striatal synapses (Maltese et al., 2018); and the structure of the nuclear envelope and accumulation of ubiquitin in the perinuclear region (Pappas et al., 2018). However, the transcriptome profile of the DYT1 model during this critical period is unknown. Such information will be informative for addressing whether and to what extent the transcriptome profile is different from the WT controls.

Here we report on our evaluation of changes in the transcriptome in DYT1 dystonia, incorporating the three points of consideration discussed above. We used a heterozygous torsinA knock-in mouse model (*Tor1a*^+/ΔE^), whose genotype is shared with DYT1 dystonia patients (*TOR1A*^+/ΔE^). We focused on samples obtained from four brain regions (the cerebral cortex, hippocampus, striatum, and cerebellum) of 2-to-3-week-old mice. We compared the transcriptome profiles of HET mice to those of WT siblings, based on DNA microarray data. In order to optimize the statistical power, we evaluated the transcriptome profiles from nine mice for each genotype, and from four brain regions per mouse, resulting in a total of 72 samples.

## 2. Materials and methods

### 2.1. Genotyping

Animal care and procedures were approved by the University of Iowa Animal Care and Use Committee, and performed in accordance with the standards set by the National Institutes of Health Guide for the Care and Use of Laboratory Animals (NIH Publications No. 80-23, revised 1996). Every effort was made to minimize suffering of the animals. On postnatal days 0-1, pups of both sexes of the ΔE-torsinA knock-in model of DYT1 dystonia (Goodchild et al., 2005) were genotyped according to a fast-genotyping procedure (within ∼5 hours, EZ Fast Tissue/Tail PCR Genotyping Kit, EZ BioResearch LLC, St, Louis, MO, USA) (Koh et al., 2015). The pups were identified long-term by tattooing the paw pads (NEO-9 Neonate Rodent Tattooing System, Animal Identification and Marking Systems Inc, Hornell, NY, USA) (Koh et al., 2015).

### 2.2. RNA preparation

We used 9 mice per genotype: 7 male and 2 female WT (*Tor1a*^+/+^) mice; and 6 male and 3 female HET (*Tor1a*^+/ΔE^) mice. They were of 2-to-3-weeks of age and obtained from two groups of littermates. The same numbers of WT and HET mice were obtained from each group. Samples of the four brain regions were obtained from each mouse (i.e. multi-regional sampling). The mice were anesthetized by intraperitoneal injection of a cocktail of ketamine (87.5 mg/kg) and xylazine (12.5 mg/kg). Mice were sacrificed one mouse at a time by decapitation, and the brain was rapidly removed and placed into ice-cold Hanks’ solution (Hanks’ Balanced Salts, without calcium chloride, magnesium sulfate and sodium bicarbonate, H2387-10X, Sigma-Aldrich, St. Louis, MO, USA) containing 4.17 mM NaHCO_3_, 10 mM HEPES, adjusted to pH 7.4 using 5 M NaOH, and adjusted 310 mOsm using sucrose. The four brain regions from each animal were dissected in the Hanks’ solution, using a binocular dissection microscope. The cerebellum included the bilateral cerebellar hemispheres and the vermis. The striatum was collected from one hemisphere (unilateral) and the whole hippocampus was obtained from the other (also unilateral). The hippocampus included both the dentate gyrus and the CA3-CA1 regions of the hippocampus proper. The cerebral cortex was bilateral, and obtained from a rostro-caudal strip in the dorso-medial portion (near the midline) of the frontal through occipital cortices.

These sizes of the brain regions were large enough that RNA from a single brain was sufficient for a single profiling experiment. To ensure consistency between samples, all dissections were carried out by the same, experienced neuroscientist who has cultured primary neurons from these regions (Koh et al., 2015). After meninges were carefully removed from the surface, the brain regions were transferred to individual, dry, ice-cold RNase-free tubes. The Hanks’ solution was removed from the tubes using a pipette, and the tubes were rapidly placed into liquid nitrogen. The dissection of each animal was completed within 9-10 min, and the four brain regions were fast-frozen within at most 11 min of decapitation. Thereafter, 1 ml of TRIzol (15596018, Thermo Fisher Scientific, Waltham, MA, USA) was added to each tube (Kim et al., 2014; Yang et al., 2016) and samples were stored at −80°C until used for RNA extraction.

Total RNA was extracted using the RNeasy Plus Mini Kit (74134, QIAGEN, Germantown, MD, USA) with modifications. The brain regions were thawed at room temperature, homogenized using nuclease-free plastic pestles and incubated on a shaker at 100 rpm at room temperature for 10-15 min. The debris were removed by centrifugation at 12,000 g for 10 min at 4°C, and the supernatant was treated with 0.2 ml chloroform at room temperature for 10-15 min in a 2-ml Phase Lock Gel tube (2302830, Eppendorf, Hauppauge, NY, USA). After centrifugation at 16,000 g for 15 min at 4°C, the clear aqueous phase was transferred to a clean tube and treated with 600 µl cold (−20°C) 70% ethanol. The sample was applied to the RNeasy spin column, centrifuged at 6,200 g for 1 min at room temperature. After the flow-through solution was discarded, the sample on the spin-column membrane was treated with 700 µl of Buffer RW1 and washed by centrifugation at 6,200 g for 1 min. After the flow-through solution was discarded, the sample on the spin-column membrane was treated with DNase I in the RNase-Free DNase Set Kit (79254, QIAGEN) at 27.3 Kunitz units / sample for 15 min at room temperature, to remove any contaminating DNA. After 350 µl Buffer RW1 was added to each spin-column membrane, it was centrifuged at 6,200 g for 15 sec, and the flow-through solution was discarded. This washing was repeated by centrifugation at 6,200 g with 500 µl mixture of 20% Buffer RPE and 80% ethanol for 15 sec, and 500 µl Buffer RPE for 2 min to eliminate the ethanol, and by centrifugation without any buffer for 1 min to eliminate any carryover of buffer. Finally, the spin-column membrane was treated with 15-30 µl RNase-free water for 5-10 min and centrifuged at 6,200 g for 1 min at room temperature to elute the RNA.

The concentration and quantity of RNA were measured using the NanoDrop Spectrophotometer (ND-1000, Thermo Fisher Scientific), and the quality was determined using the RNA PICO chip of Agilent 2100 Bioanalyzer (Agilent, Santa Clara, CA, USA). RNA Integrity Number (RIN) was used to assess the overall quality of the prepared RNA (Imbeaud et al., 2005; Schroeder et al., 2006). For all samples, the RIN value was 9.05 ± 0.04 (mean ± sem, n = 72 samples), with a minimum and maximum value of 7.9 and 9.8. These values met the criteria for our subsequent analyses. RIN values were not statistically different between any pair of brain regions (cerebral cortex, 9.01 ± 0.07; hippocampus, 9.09 ± 0.04; striatum, 9.11 ± 0.07; cerebellum 8.98 ± 0.11; n = 18 samples for each group) or between genotypes (HET 9.08 ± 0.04; WT 9.02 ± 0.06; n = 36 samples for each group), with the resulting p>0.29 for any comparison (unpaired *t*-test).

### 2.3. cDNA, cRNA preparation and DNA microarray sample processing

The transcriptome was analyzed using the Illumina Whole-Genome Gene Expression microarray system (Illumina, San Diego, CA, USA) in a one-color labeling scheme. RNA sample preparation for hybridization and the subsequent hybridization to the Illumina BeadChips were performed at the Iowa Institute of Human Genetics, using the manufacturer’s recommended protocol. Briefly, 100 ng of total RNA containing Poly(A) RNA were reverse transcribed into cDNA, converted to double-stranded cDNA, and *in vitro* transcribed into amplified and biotinylated cRNA (Biotin-cRNA), using the Epicentre TargetAmp-Nano Labeling Kit for Illumina Expression BeadChip (TAN07924, Illumina) according to the manufacturer’s recommended protocol. The amplified Biotin-cRNA product was purified using a QIAGEN RNeasy MinElute Cleanup column (74204, QIAGEN) according to modifications from Epicentre. 750 ng of this product were mixed with Illumina hybridization buffer, loaded into each array of MouseRef-8 v2.0 Expression BeadChips (BD-202-0202, Illumina), and incubated at 58°C for 17 h, with rocking, in the Illumina Hybridization Oven. Following hybridization, the arrays were washed, blocked, and stained with streptavidin-Cy3 (Amersham-GE Healthcare, Piscataway, NJ, USA) according to the Illumina Whole-Genome Gene Expression Direct Hybridization Assay protocol. BeadChips were scanned using the Illumina iScan System, and data were collected using the GenomeStudio Software v2011.1 (Illumina), with a 16-bit dynamic range of 0-65,535 fluorescence intensity units.

Eight samples were processed in a single MouseRef-8 v2.0 Expression BeadChip with an 8-array format. Each BeadChip was used to analyze 4 brain regions of a single WT mouse and 4 brain regions of a single HET mouse. Using this genotype-pairing design, we analyzed 18 brains (i.e. 72 brain regions) in a total of 9 BeadChips. In order to avoid systematic bias in the microarray data and to minimize inter-chip variations (batch effects), the mice were paired after the brain collection dates, RNA extraction dates and sexes were randomized. An additional (i.e. tenth) BeadChip was used for technical replicates. For this purpose, RNA samples from 8 out of 9 HET mice were randomly selected, and for each mouse, 1 of the 4 brain regions was randomly assigned (for a total of 2 samples for each brain region from HET mice) and loaded into the separate BeadChip. Thus, 8 of the 72 original RNA samples were analyzed twice using the microarray method.

### 2.4. Microarray analysis

We analyzed the transcriptome information of the same set of 72 samples = (4 brain regions / mouse) × (9 mice / genotype) × (2 genotypes), excluding the technical replicates mentioned above. The information from these 72 samples was grouped in three different ways, for comparing the transcriptome profiles in: 1) distinct brain regions (18 samples / brain region), 2) the same brain regions in distinct genotypes (9 samples / brain region / genotype; comparison within tissue), and 3) distinct genotypes in all brain regions (36 samples / genotype; global comparison).

Microarray data were imported and normalized using the “beadarray” (2.28.0) package in R (3.4.3), running under OS X 10.11. Quantile normalization was performed followed by a log_2_ transformation (Bolstad et al., 2003). The 24,482 probes for 16,886 genes with unique Ensembl IDs were used for differential expression modeling. Linear modeling of the transformed data was performed using Linear Models for Microarray Data (Limma, 3.34.9) with the “eBayes” empirical Bayes moderation of standard errors (Smyth, 2004; Smyth, 2005). For testing genotype effects in the same brain regions, a model was created that was conditioned on a combined tissue and genotype factor with eight levels, according to best practices recommended by the “DESeq2” package (Love et al., 2014). For testing genotype effects in global comparison, a model was created that accounted for genotype and tissue as independent factors. We did not attempt to account for genotype-tissue factor interactions in this study. Testing on sex showed that it did not contribute significantly to the variation observed. Significance values (“P” values) and false discovery rate (FDR)-adjusted P values were obtained for each group-wise comparison of interest within each model. Only differences in expression levels with the adjusted P value <0.01 were considered indicative of differential expression. Fold change was calculated as the ratio of the individual gene signals for one group to those of another group (e.g. x / y for group X vs. group Y). The fold change with respect to genotypes was always calculated as HET/WT; thus a fold change >1, or log_2_(fold change) >0, indicates that a gene was upregulated in HET samples in comparison to WT samples.

The raw microarray data are available at: (*Currently under processing; URL is to be attached here in the final manuscript*). The full analysis script written in R is available at: https://github.com/mchimenti/harata_dyt1_analysis_scripts.

The technical replicates were used to evaluate technical reproducibility and the possibility of batch effects in the microarray results (Fig. S1). The measured fluorescence intensity for individual probes on a log_2_ scale was used to generate correlation plots for the eight pairs of original samples and their technical replicates. Linear regression lines were drawn without any constraints. The correlations were high, with the Spearman correlation R values ranging from 0.943 and 0.991 (0.966 ± 0.007, mean ± sem, n = 8 samples). This result reinforces the validity of our approach and data. Slight variation was visible in each pair, i.e. each measurement: this unavoidable noise was suppressed by including a high number of biological replicates in our analyses (n = 9 mice for each brain region).

Principal component analysis (PCA) was performed using the “prcomp” package from the base ‘stats’ R library. Dendrograms and heat maps were generated by hierarchical clustering using the Euclidian distances of all genes as a metric. Pathway analysis was performed using the Ingenuity Pathway Analysis (IPA) (content version 47547484, QIAGEN, Germantown, MD, USA).

## 3. Results

### 3.1. Overall patterns of transcriptome profiles

Two methods were used to visualize the relatedness of transcriptome profiles in all 72 samples (Fig. 1). Principal component analysis (PCA) was first used to evaluate all probes (Fig. 1A-C). Samples with similar gene expression patterns would be plotted in the 2-dimensional neighborhood. Samples from the cerebellum and striatum formed independent clusters, whereas those of the cerebral cortex and hippocampus formed a merged cluster (Fig. 1A). The samples did not cluster by either genotype (HET and WT, Fig. 1B) or sex (males and females, Fig. 1C).

**Fig. 1.**
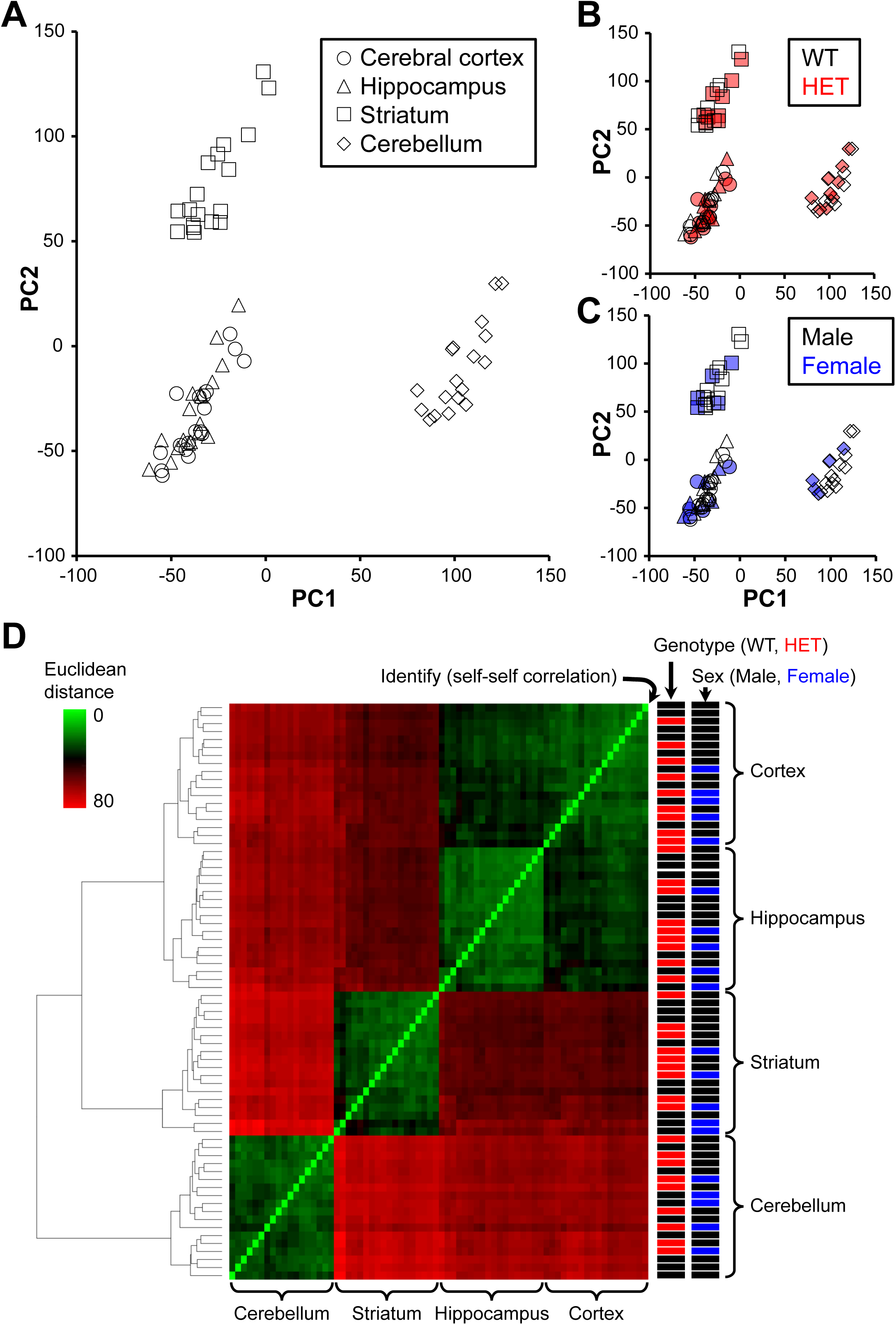
Overall patterns of gene expression in four brain regions of wild-type (WT) and heterozygous (HET, *Tor1a*^+/ΔE^) mice. (A-C) Principal component analysis, with the individual symbols representing individual samples. Proximity of symbols indicates transcriptional similarity between two samples. PC1 and PC2 represent the first and second principal component. (A) The symbols for the four brain regions are indicated in the legends. (B) Genotypes were used as the sample labels. (C) Sexes were used as the sample labels. (D) Hierarchical clustering. Dendrogram branching shows hierarchical clustering of all samples. Heat map represents correlation between two samples. The x- and y-axes list all samples in the same order. The brightest green symbols along the diagonal line represent identity (i.e. Euclidean distance = 0). Attribute bars at right show the properties of the samples: genotype (WT, black; HET, red) and sex (male, black; female, blue). In all panels of this figure, the total number of samples was 72 samples = (9 mice / genotype) × (4 brain regions / mouse) × (2 genotypes).

Hierarchical cluster analysis was used to generate a second representation of sample relatedness (Fig. 1D). In this analysis, dendrogram branching and heat map of correlations reveal the degree of transcriptional similarity between any two samples. The samples from the cerebellum and striatum formed independent clusters, and samples from the cerebral cortex and hippocampus formed related but separate clusters. Thus the samples were clearly clustered according to the brain regions. Again, the samples did not cluster by genotype or sex, either within a brain region or across brain regions (attribute bars at far right, Fig. 1D).

These results show that, of the three attributes examined (brain region, genotype, and sex), the samples clustered only by brain region. This indicates that the overall transcriptome patterns in our samples were determined largely by the brain regions.

### 3.2. Differential expression of genes between different brain regions

Differences in the transcriptome were next evaluated at the level of individual genes. Pairwise comparisons among the four brain regions were performed and the results are illustrated by volcano plots (Fig. 2). For each pair, genes were considered to be differentially expressed with the adjusted P value of <0.01 (blue symbols in Fig. 2). The number of differentially expressed genes was highest for the comparison between the cerebellum vs. cerebral cortex (6005 probes for 4907 genes, Fig. 2C), and it was lowest for the comparison between the hippocampus vs. cerebral cortex (976 probes for 855 genes, Fig. 2A). The fold-change was highest for the comparison between cerebellum vs. cerebral cortex (6.57 on a log_2_ scale; 94.82 on a linear scale; Fig. 2C), and smallest for the comparison between the cerebellum vs. hippocampus (−5.12 on a log_2_ scale; 0.03 on a linear scale; Fig. 2E). These results indicate strong differences in the brain region transcriptome (Figs. 1, 2), as had been reported previously for WT human (Kang et al., 2011) and rat (Stansberg et al., 2007) brains.

**Fig. 2.**
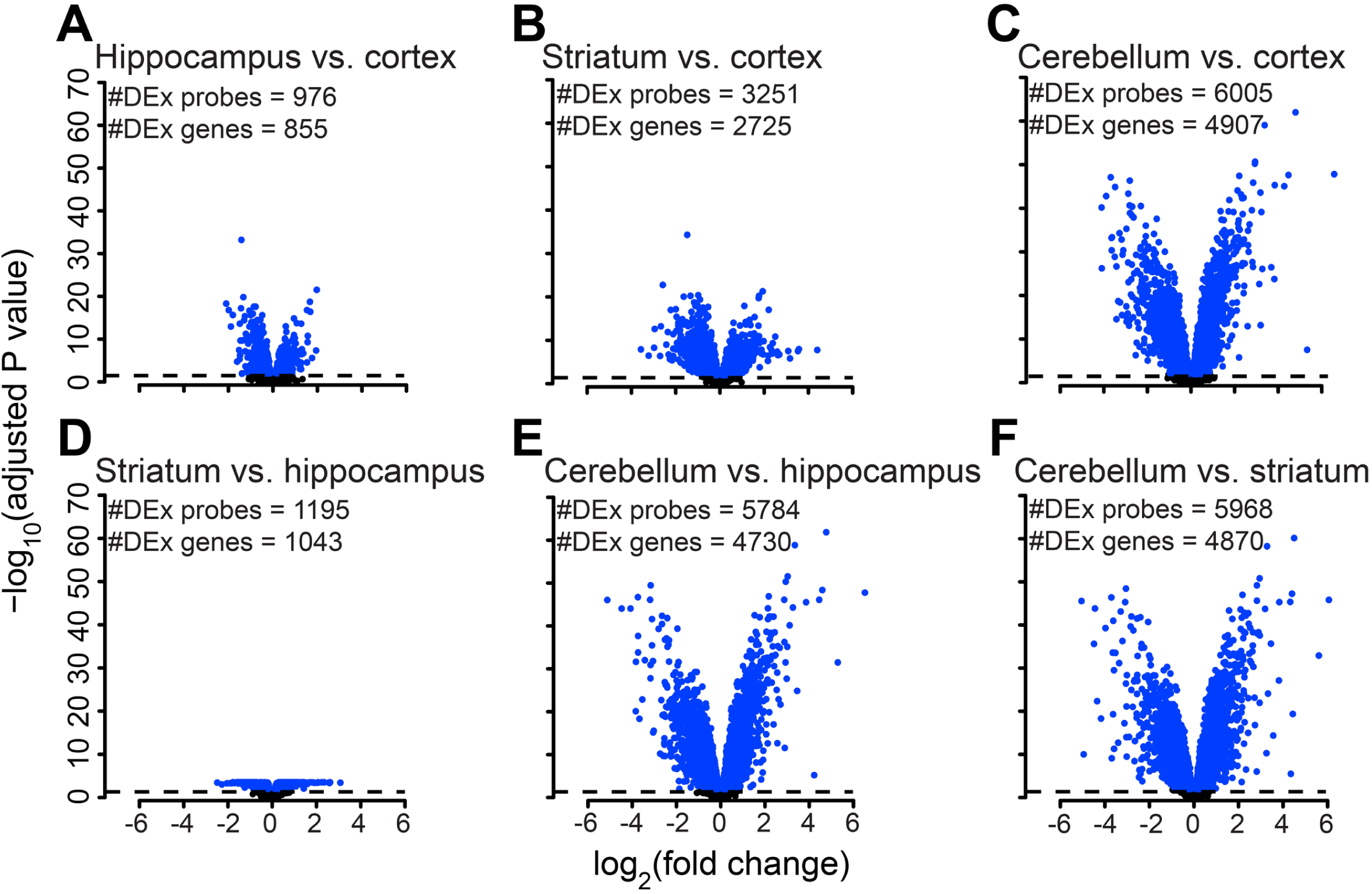
Differential expression of multiple genes by brain region. Transcriptomes in the four brain regions compared in a pairwise manner, with the numbers of differentially expressed probes for specific genes: (A) hippocampus vs. cerebral cortex (976 probes for 855 genes), (B) striatum vs. cerebral cortex (3251 probes for 2725 genes), (C) cerebellum vs. cerebral cortex (6005 probes for 4907 genes), (D) striatum vs. hippocampus (1195 probes for 1043 genes), (E) cerebellum vs. hippocampus (5784 probes for 4730 genes), and (F) cerebellum vs. striatum (5968 probes for 4870 genes). In these volcano plots for gene-level comparisons, the x- and y-axes represent the magnitude of the change (log_2_(fold change)) and the measure of statistical significance (−log_10_(adjusted P value)) for individual probes, respectively. Blue filled circles represent differentially expressed probes, identified at the adjusted P value <0.01. #DEx, number of differentially expressed probes or genes. Number of samples was 18 mice / brain region = (9 mice / brain region / genotype) × (2 genotypes).

### 3.3. Differential expression of genes between genotypes within specific brain regions

Genotypic differences in the transcriptome within each brain region are illustrated by volcano plots (Fig. 3). Note that the x- and y-axis scales necessary to show all the genes are very different from those in comparisons across brain regions (Fig. 2), indicating that the overall changes in HET brain were modest if present. The only differentially expressed gene in any of these samples was *Gprin3* [G protein-regulated inducer of neurite outgrowth (Gprin) family member 3], in the striatum. Its adjusted P value was 7.21 × 10^−3^, and log_2_(fold change) was 0.29, i.e. with only a 1.22-fold upregulation in HET brain.

**Fig. 3.**
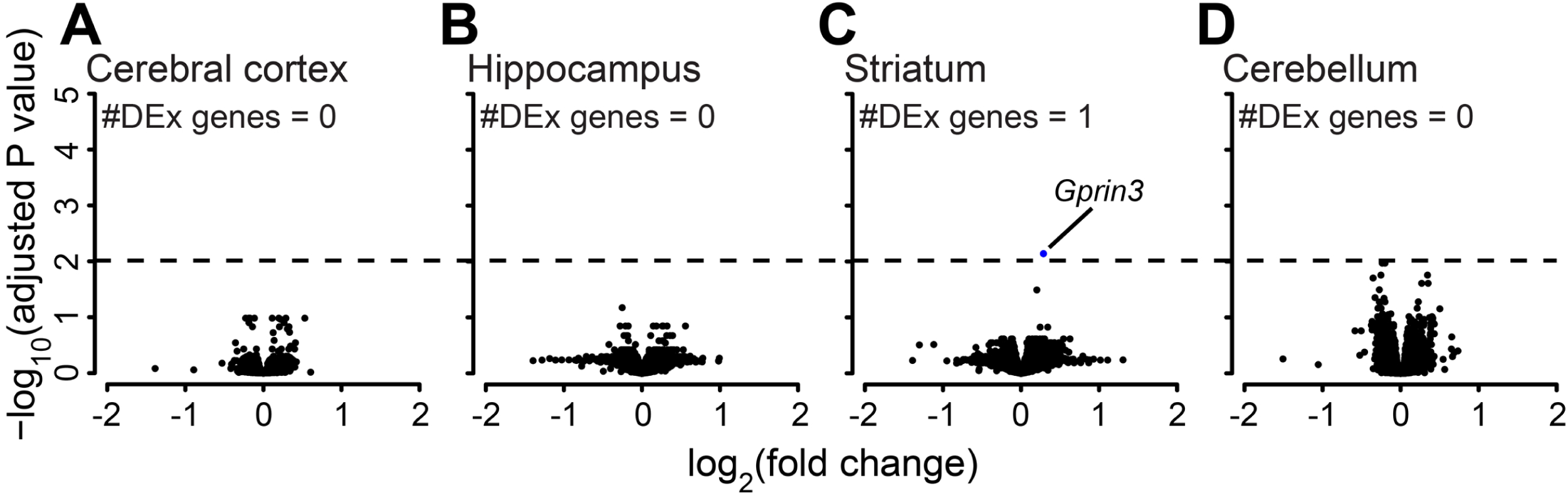
Gene-level comparison of transcriptomes between mice of different genotypes, by brain region. Comparison of transcriptomes of HET vs. WT brains for: (A) cerebral cortex, (B) hippocampus, (C) striatum, and (D) cerebellum. Volcano plots are drawn as in **Fig. 2**. Number of samples was 9 mice / brain region / genotype.

### 3.4. Differential expression of genes between genotypes in combined brain samples

Genotypic differences in the global transcriptome were also assessed for the combined samples. This approach increased the statistical power by increasing the number of analyzed samples to n = 36 per genotype. The volcano plot shows the data for 308 differentially expressed probes for 301 genes (Fig. 4A). Among the 301 differentially expressed genes, seven genes were represented by two probes each. It was reassuring that these two probes showed the same direction and the similar degree of changes for each gene: log_2_(fold change) = 0.10, 0.14 for *Brcc3*; 0.23, 0.29 for *Eif4g2*; 0.17, 0.22 for *Etfa*; 0.17, 0.44 for *Xist*; −0.25, −0.27 for *Ank1*; −0.12, −0.15 for *Smim26*; −0.13, −0.14 for *Zfp64*. A part of the volcano plot is magnified to show the top 16 highly significant genes with the adjusted P value <0.0001 (i.e. 10^−4^) (Fig. 4B).

**Fig. 4.**
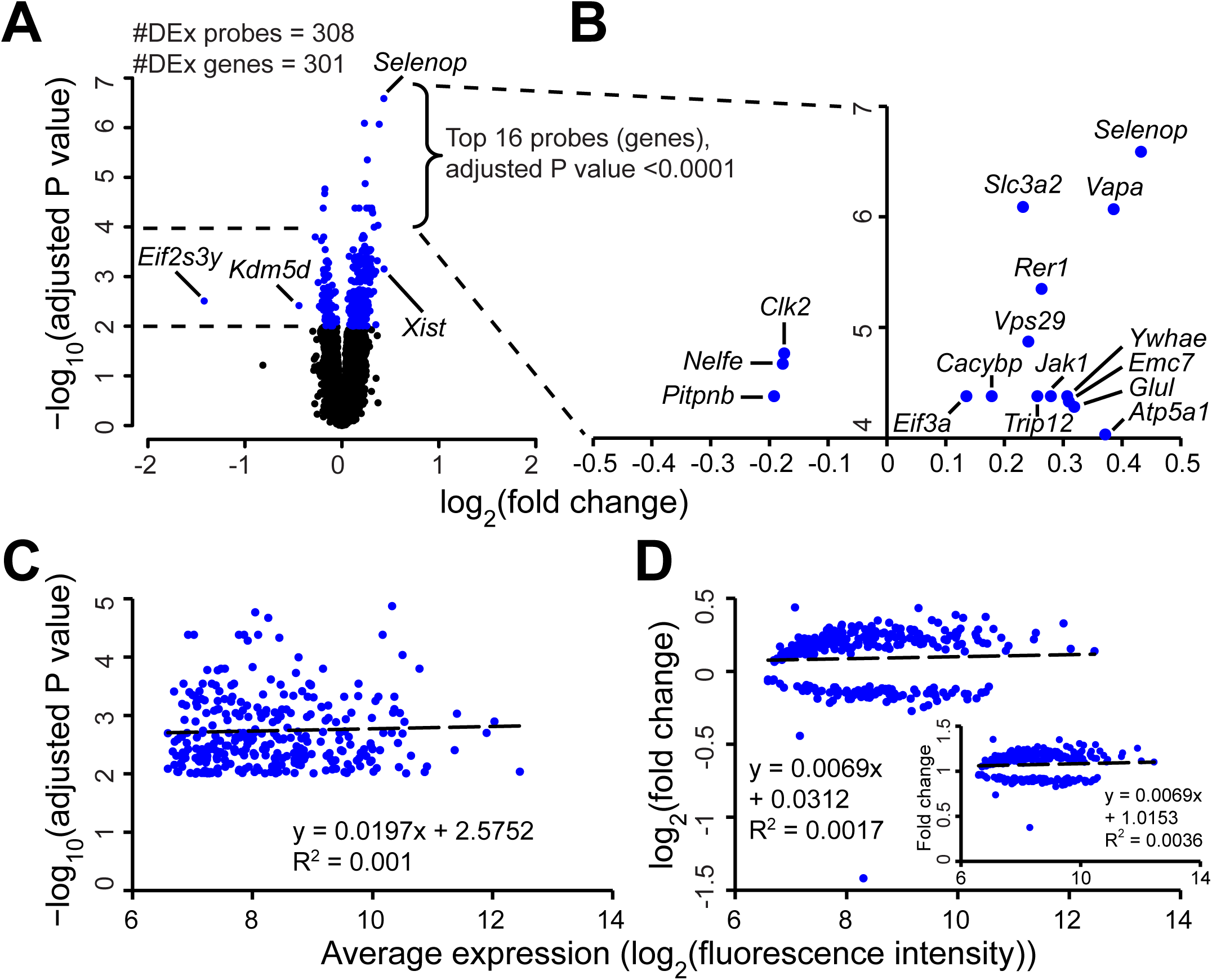
Gene-level comparison of transcriptome between different genotypes in a global, 4-brain-region combination. Information for samples of a single genotype (HET or WT) was combined irrespective of brain region. Number of samples was 36 samples / genotype = (9 mice / genotype) × (4 brain regions / mouse). (A) Volcano plot showing differentially expressed probes at adjusted P value <0.01. The names of the genes with the most extreme fold-changes are also shown. (B) The same Volcano plot but showing only the top 16 highly significant genes with adjusted P value <0.0001 (i.e. 10^−4^), and their names. (C,D). Plots of correlation between the average expression levels for all 72 samples (log_2_(fluorescence intensity)) and either the statistical significance (−log_10_(adjusted P value), C) or fold change (log_2_ scale, D). Blue filled circles in all panels represent differentially expressed probes, identified at the adjusted P value <0.01.

These plots illustrate that all differentially expressed genes were only slightly up- or down-regulated in terms of fold change (Fig. 4A,B). The two genes with the most significant down-regulation in HET samples were *Eif2s3y* (eukaryotic translation initiation factor 2, subunit 3, structural gene Y-linked) (log_2_(fold change) = −1.42 or fold change = 0.37) and *Kdm5d* (lysine demethylase 5D) (log_2_(fold change) = −0.44 or fold change = 0.74). The two genes with the most significant up-regulation in HET samples were: *Xist* (inactive X specific transcripts) (log_2_(fold change) = 0.44 or fold change = 1.35) and *Selenop* (selenoprotein P) (log_2_(fold change) = 0.43 or fold change = 1.35). Thus, the majority of differentially expressed genes showed small changes: −0.30< log_2_(fold change) <0.40, or 0.81< fold change <1.32 (i.e., from an ∼19% decrease to a 32% increase). The list of all differentially expressed probes detected in our analysis is shown in the Supplementary Table S1.

Among all differentially expressed genes, the absolute expression level (averaged over all 72 samples) did not correlate with either statistical significance (Fig. 4C) or fold change (Fig. 4D). This result indicates that the identification of differentially expressed genes was not affected by expression levels.

### 3.5. Expression levels of individual genes

The global analysis identified differentially expressed genes between the HET and WT mice (Fig. 4), in spite of the facts that 1) such differences were not detectable in individual brain regions (Fig. 3), and 2) differences in gene expression between these brain regions were highly significant (Figs. 1-2). These results suggested that the genotype-related differences within each brain region were small, but they were detectable by increasing the number of analyzed samples, and treating the genotype and brain region as independent analysis factors.

Figure 5 shows the expression levels of the identified genes, in the 4 brain regions and in the all-combined samples (Fig. 5). In the box plots, the expression levels (i.e. raw fluorescence intensities) of the 16 genes with the highest significance were compared (Fig. 4B, 5A), as were the expression levels of the two genes with the highest fold change (Fig. 4A, 5B). The majority of these 18 genes show consistent, directional changes. However, some genes show bi-directional changes across regions, revealing a genotype-factor interaction that was not accounted for in this analysis.

**Fig. 5.**
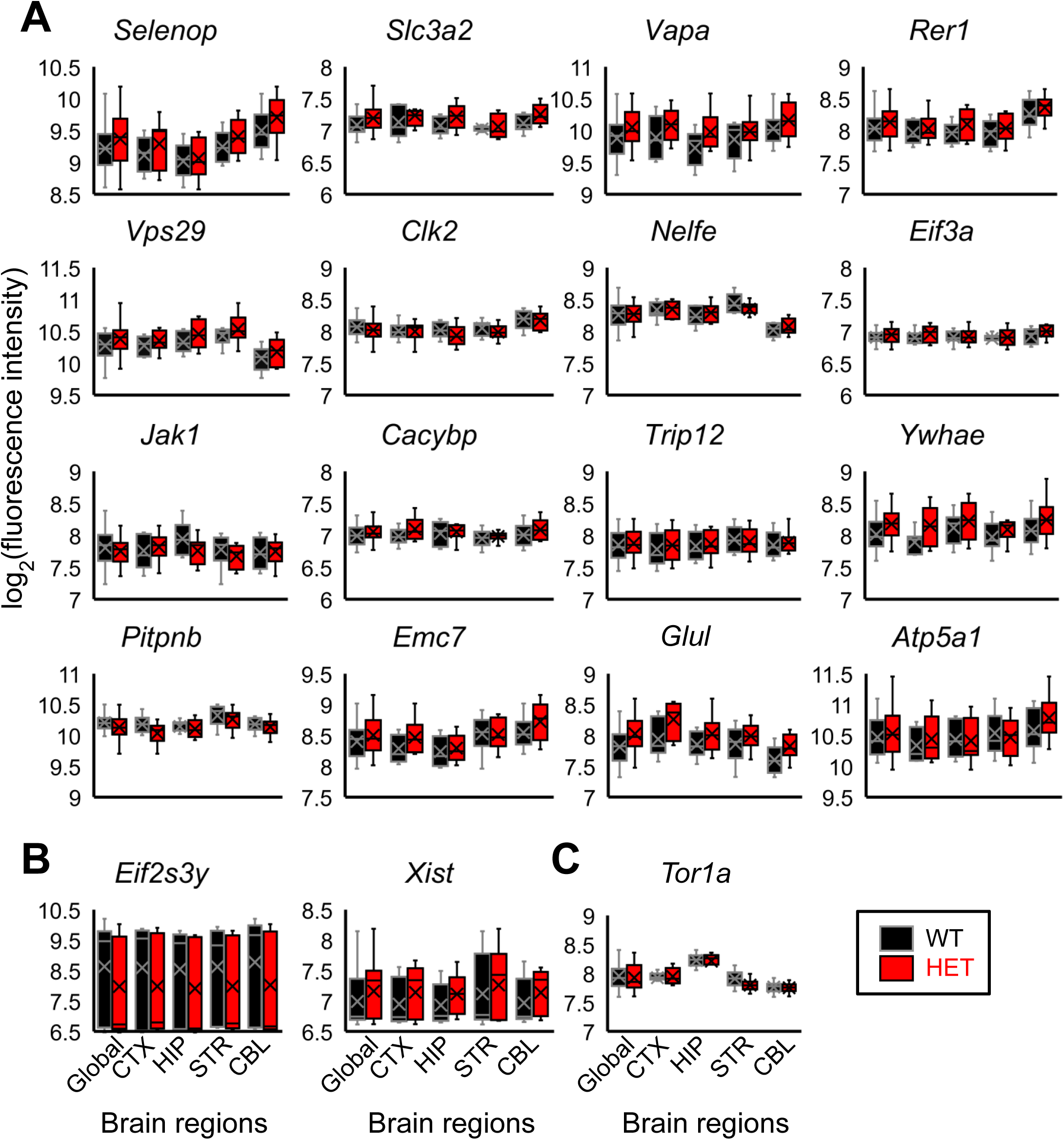
Global comparison of levels of expression for individual genes in mice of different genotypes. Box plots are shown for: (A) 16 highly significant genes with the adjusted P values of <0.0001 (gene names indicated in Fig. 4B), (B) differentially expressed genes with the minimal (left) and maximal (right) log_2_(fold change) values (gene names indicated in Fig. 4A), and (C) *Tor1a* gene, which is mutated in the current mouse model. In the “global” analysis, the number of samples was 36 samples / genotype = (9 mice / genotype) × (4 brain regions / mouse), as in Fig. 4. In each brain region, the number of samples was 9 mice / brain region / genotype, as in Fig. 3. Abbreviations for brain regions: CTX, cerebral cortex; HIP, hippocampus; STR, striatum; CBL, cerebellum. A box indicates the levels of the 25^th^, 50^th^ (median) and 75^th^ percentile. “x” represents the mean value. The vertical bars extending from a box indicate the minimum and maximum values, excluding the outliers whose values lie above the 75^th^ percentile or below the 25^th^ percentile by more than 1.5 × (75 percentile − 25 percentile).

Among the 18 genes, five genes (*Selenop*, *Rer1*, *Vps29*, *Clk2*, *Nelfe*) were found in at least one brain regional comparison at the adjusted P value of <0.01 (Fig. 2, Supplementary Table S1). All of five them were found in relation to the cerebellum (i.e. cerebellum vs. cerebral cortex or hippocampus), implying the high impact of the cerebellar expression of these genes on the global transcriptome changes. The box plot of *Tor1a* is also included for comparison (Fig. 5C). *Tor1a* was not differentially expressed, and the box plot supports this lack of change. The finding of unchanged *Tor1a* expression is consistent with a report of its mRNA levels in the HET brain (Goodchild et al., 2005).

### 3.6. Interpretation of differentially expressed genes

Pathway analysis of the 301 differentially expressed genes identified multiple canonical pathways (Fig. 6). The top five were: mitochondrial dysfunction, oxidative phosphorylation, protein ubiquitination pathway, regulation of eIF4 and p70S6K signaling, and cell cycle: G2/M DNA damage checkpoint regulation, in the decreasing order of significance. Additional pathways are directly involved in neuronal excitability control, e.g. acetyl-CoA biosynthesis (precursor of acetylcholine), α-adrenergic signaling, protein kinase A signaling, and axonal guidance signaling.

**Fig. 6.**
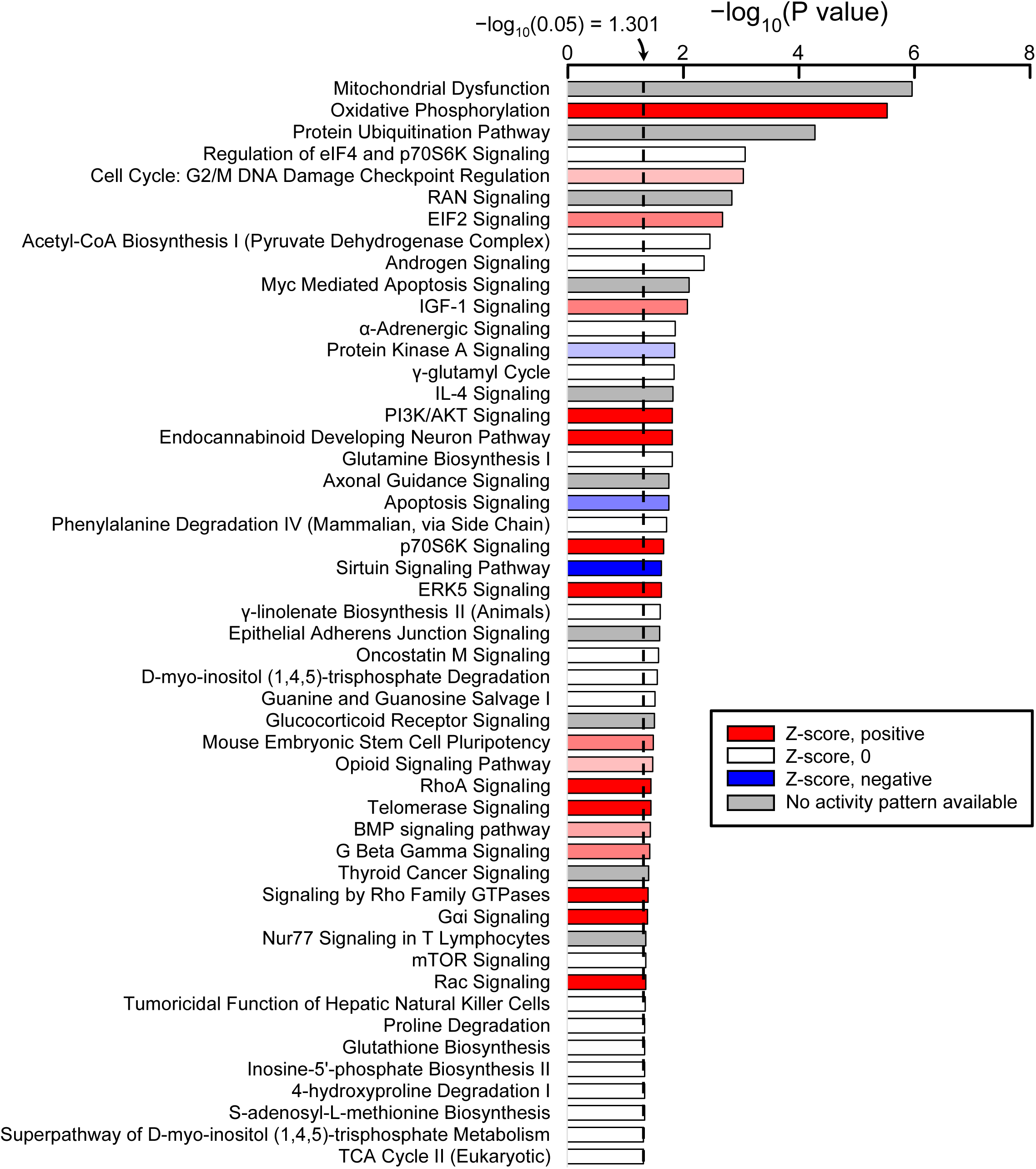
Pathway analysis based on differentially expressed genes detected in global analysis. X-axis represents the statistical significance (−log_10_(P value)), with the broken line indicating the 0.05 threshold for this analysis. The color of a horizontal bar indicates the z-score.

Manual curation among 90 highly significant genes (adjusted P value <0.001, i.e. 10^−3^) identified the following gene groups of our interest, based on domain knowledge: mitochondrial function (*Atp5a1*, *Atp5e*, *Atp5l*, *Cycs*, *Dld*, *Etfa*, *Higd1a*, *Mrpl20*, *Oxct1*, *Pdhb*, *Ppox*, *Tomm70a, Vdac1*), ubiquitination/deubiquitination proteasome pathway (*Brca1*, *Btbd6*, *Cacybp*, *Huwe1*, *Psmc6*, *Trip12*, *Ubac1*, *Usp14, Yod1*), eukaryotic translation initiation (*Eif2s3x*, *Eif3a*, *Eif4g2, Pabpc1*, *Rps3*), intracellular trafficking (*Bcap31*, *Fig4*, *Ift52*, *Ktn1*, *Pitpnb*, *Rab5a*, *Rab10*, *Rer1*, *Tmed2*, *Vapa*, *Vps29*), cytoskeletal regulator (*Ank1*, *Cfl2*, *Kif21b*, *Ktn1*, *Nphp4*, *Sept7*), intracellular signaling (*Glul*, *Impa1*, *Jak1*, *Lpgat1*, *Ntrk3*, *Ppp6r3*, *Ptprr*, *Rack1*, *Rap1b*, *Rraga*, *Rsu1*, *Ywhae*), extracellular signaling (*Adam9*, *Hapln1*, *Nlgn1*, *Tjp1*), nuclear pore transport (*Hnrnpa2b1*, *Kpna3*), heat shock protein (*Hspa5*), autophagy (*Wdr45*), AAA+ ATPase (*Pex6*), and neuronal excitability control (*Gng5*, *Nlgn1*, *P2ry12*).

## 4. Discussion

Our study of the transcriptome in the DYT1 dystonia mouse brain is distinguished from previous studies in mammalian models (**Table 1**) by the following four points collectively: 1) we used the *Tor1a*^+/ΔE^ mice which share the same mutant state as DYT1 dystonia patients, 2) we analyzed multiple brain regions (the cerebral cortex, hippocampus, and cerebellum as well as the more commonly studied striatum), 3) we assessed the critical postnatal window during which the mutated *Tor1a* gene affects the brain, and 4) we used a high “n” to achieve robust statistical power (n = 9 for comparisons of the transcriptome across brain regions, and n = 36 for comparisons of global transcription). These experimental conditions were chosen for their likelihood of yielding meaningful results. It is notable that genotype-specific differences in transcriptome were not detected in any individual brain region, except for a single gene in striatum, yet such differences became apparent when the data from the four brain regions were combined to simulate a whole-brain comparison. These results suggest that the *Tor1a*^+/ΔE^ genotype causes changes in the brain transcriptome, but the change is very subtle, with most changes being consistent in direction across the brain regions.

### 4.1. Validity of the current data

Two observations support the appropriateness and validity of the experimental methods used in the current study for identifying transcriptome changes. First, the transcriptome profiles in the analyzed brain regions were distinct (Figs. 2, 3), consistent with previous reports for WT brains of both rats (Stansberg et al., 2007) and humans (Kang et al., 2011). This validates the consistency and accuracy of our dissection of brain regions and our analyses of the transcriptome data.

The second observation that validates our approach is that slight variations inherent to the experimental system, as visualized in technical replicates (Fig. 1), were overcome when a large number of samples was compared. The DNA microarray method can provide quantitative analytical data (Liu et al., 2014), especially for low-copy-number genes, without being affected by the “depth” of sequencing. It has some disadvantages relative to RNA sequencing, e.g. less dynamic range, inability to discover new genes because the detectable genes need to be pre-determined and annotated, and inefficiency in estimating exon-specific and single-base-pair events (Dillman et al., 2013; Liu et al., 2014; Shendure, 2008). However, the results of this study are not significantly affected by such disadvantages, because our genotypic comparison is limited to the well-annotated genes, and evaluation of the results is focused on detecting the presence of small changes (addressing whether any transcriptome change occurs or not) rather than large changes (addressing, e.g., which brain region shows the largest fold-changes).

### 4.2. Implications of genotype-dependent differences in gene expression

Four points merit discussion regarding the transcriptome changes obtained in this work. The first is the fact that transcriptome changes were detectable in the critical postnatal window. This is significant, in light of a general notion that young age (as evaluated in the current study) is the time period when gene expression profiles demonstrate minimal differences, resulting in a so-called temporal “cup-shaped or hourglass-like” pattern, e.g. in human and non-human primate brains (Li et al., 2018; Pletikos et al., 2014; Zhu et al., 2018).

The second point is that the fold-change values for differentially expressed genes were small in the global analysis. The majority of the differentially expressed genes showed between 19% decrease to 32% increase, and a large sample number (n = 36 samples / genotype) was required to detect them. These small changes likely account for the challenge in detecting differentially expressed genes in individual brain areas (Fig. 4). Our results indicate that the *Tor1a*^+/ΔE^ genotype impacts the brain transcriptome, yet the effects were not drastic in amplitude, at least during the period studied (2 to 3 weeks after birth).

The third point is the nature of the pathways and individual genes that are affected in *Tor1a*^+/ΔE^ mice. Several of these had been implicated previously, for example, the eukaryotic translation initiation (Beauvais et al., 2018; Beauvais et al., 2019; Rittiner et al., 2016), protein ubiquitination pathway (Gordon et al., 2012; Nery et al., 2011) and cell cycle (Ledoux et al., 2013). Others were of particular interest, such as the mitochondrial function, intracellular trafficking and axonal guidance: they are involved in regulating neuronal functions including excitability. It is expected that these gene transcriptome changes underlie key aspects of dystonia pathophysiology, since small changes over a wide variety of gene networks can impact neuronal excitability (Noebels, 2015).

The fourth point relates to the direction of changes in the 301 differentially regulated genes. 209 of them (69%) were upregulated. In retrospect, our finding was consistent with the approaches in the previous report (Rittiner et al., 2016). Gene knockdown by an RNA interference strategy was effective in screening for genes that correct the mislocalization of EGFP-tagged human ΔE-torsinA in HEK cells. Among the 93 genes identified in that report, two were identified in our global analysis: *Atp5a1* (ATP synthase, H+ transporting, mitochondrial F1 complex, alpha subunit 1; human ortholog *ATP5F1A*; adjusted P value = 9.27 × 10^−5^) involved in ATP synthesis in mitochondria, and *Trappc3* (trafficking protein particle complex 3; adjusted P value = 6.79 × 10^−3^) involved in vesicular transport from the endoplasmic reticulum to Golgi apparatus. Both genes were upregulated in HET brains in our result, consistent with expectations based on the previous work (Rittiner et al., 2016).

### 4.3. Implications for future work on genotype-dependent differences in gene expression in specific brain regions

There are at least two implications of the fact that drastic changes were not observed in the brain regional transcriptomes of *Tor1a*^+/ΔE^ mice.

A positive change in the transcriptome of a particular brain region could have been undetectable because of heterogeneity within each region. It may be that subtle and unique changes are present in specific cellular and subcellular components, in addition to the universal change across brain regions. Any brain region is composed of multiple cell types, e.g. neurons, glial cells, endothelial cells. In addition, within a single brain region, further heterogeneity arises from the presence of somata and dendrites of multiple types of neurons, and from the fact that these neurons receive afferent axons and nerve terminals from neurons outside the region. Since the subcellular components, i.e. dendrites and axons, contain mRNA and the translational machineries separately from those in soma (Biever et al., 2019), a true picture of brain regional transcriptome can be inevitably complex. In future studies, it will be useful to take heterogeneity into account and to use a more individualized approach for analyzing the transcriptome, e.g. at the level of specific cell types or single cells.

It is possible that transcriptome changes in individual brain regions are extremely low and therefore undetectable. However, a lack of large transcriptome changes does not necessarily indicate a lack of functional change. For example, lateralization of brain function has been noted in the human neocortex in spite of a lack of detectable hemispheric differences in the transcriptome (Hawrylycz et al., 2012; Pletikos et al., 2014). In future, it would be desirable to use RNA sequencing, which is more sensitive than the DNA microarray method, and to evaluate complexity of the transcriptome at levels beyond gene abundance. For example, multiple splice variants could lead to proteomic diversity, and the RNA editing by adenosine deaminase could lead to modification of the amino acid sequence or to differences in splicing or nuclear retention of the transcript (Dillman et al., 2013; Zhao et al., 2014). Furthermore, even in the absence of drastic changes in the transcriptome, translational regulation could be strongly affected (Martin et al., 2009). Thus, protein expression analysis might be also informative, as has been demonstrated by proteomics approaches (Beauvais et al., 2016) and the analyses of individual proteins (Vanni et al., 2015).

## 5. Conclusion

This study demonstrated the presence of transcriptome changes throughout multiple brain regions of *Tor1a*^+/ΔE^ mice. This is important for dystonia, a network disorder involving multiple brain regions. Although the changes were mild, the detected gene pathways suggest potential directions to some focused research, especially at the cellular level. This study also has some implications for future studies, involving, e.g. the brain regional transcriptome and proteome.

## ABBREVIATIONS

AAA+: ATPases associated with various cellular activities
CBL: cerebellum
CTX: cerebral cortex
ΔE- torsinA: mutated form of torsinA
#DEx: number of differentially expressed probes or genes
FC: fold change
HEK: human embryonic kidney cells
HET: heterozygous ΔE-torsinA knock-in mouse
HIP: hippocampus
PC: pheochromocytoma
PCA: principal component analysis
PCn: n-th principal component
STR: striatum
WT: wild-type

## Acknowledgments

The authors thank Drs. Elizabeth Scholl, Amy Lee and Stefan Strack (The University of Iowa) for instructions and comments on the RNA extraction protocol, and Drs. Mark Stamnes and Kai Wang (The University of Iowa) for discussions. This work was supported by the US Department of Defense (W81XWH-14-1-0301 to NCH) and Dystonia Medical Research Foundation (to NCH). Declarations of interest: none.

## Supplementary Table Legend

**Table S1.** The Excel file contains information about the differentially expressed probes in seven sheets. The first six sheets (e.g. “HIP vs CTX”) show the brain regional comparisons, irrespective of genotypes; The pairwise comparisons correspond to the results shown in **Fig. 2**. Brain regions were labeled as: CTX, cerebral cortex; HIP, hippocampus; STR, striatum; CBL, cerebellum. The last sheet (“Global-HET vs WT”) shows a global comparison after the 4-brain-region combination; The comparison corresponds to the results shown in **Figs. 4-6**. Each sheet shows: 1) gene symbol (SYMBOL), 2) log_2_(fold change) (log2(FC)), 3) average fluorescence intensity of all 72 samples probes in a log_2_ scale (AveExpr), and 4) adjusted P value (adj.P.Val) of each probe.

**Fig. S1.**
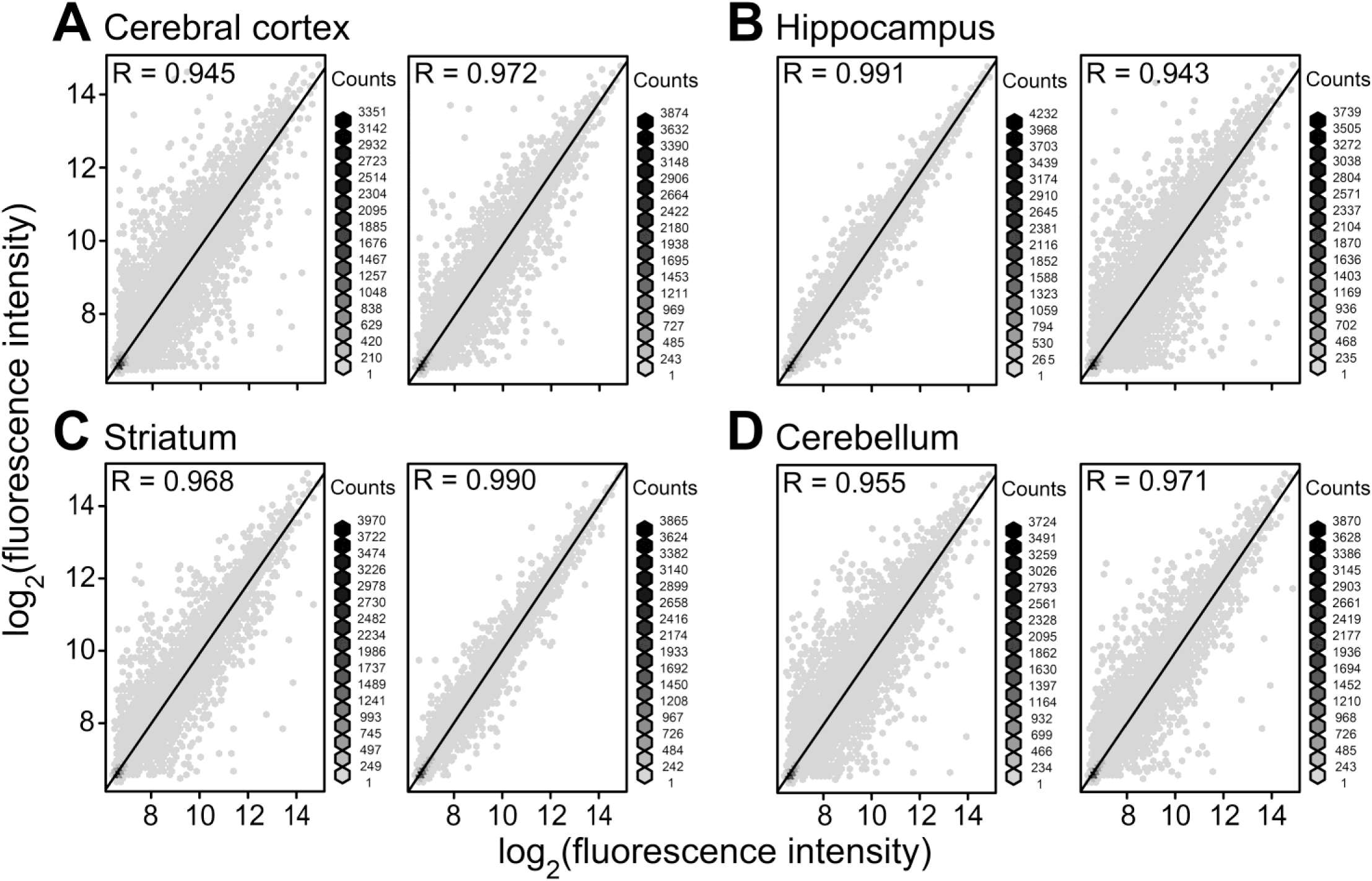
Reproducibility of the analysis system, as evaluated by technical replicates. Eight out of the 72 original RNA samples were analyzed twice by the microarray method, and the expression of individual genes is shown in the correlation plots. All samples were obtained from heterozygous (HET, *Tor1a*^+/ΔE^) mice. The brain regions and the mouse identification numbers (serial #’s for nine mice) were: **(A)** cerebral cortex with #2 (male) and #4 (male), **(B)** hippocampus with #1(male) and #8 (female), **(C)** striatum with #6 (female) and #9 (male), **(D)** cerebellum with #5 (female) and #7 (male). For the presentation, overlapping symbols were avoided by using a hexbin format. A plotting window is represented as a collection of non-overlapping hexagons (hexagonal bin or hexbin), and the number of genes per hexbin is shown coded in grayscale. All axes of the correlation plots represent the measured fluorescence intensity of individual probes in a log_2_ scale. Spearman correlation R values are shown.

